# Visualization and normalization of drift effect across batches in metabolome-wide association studies

**DOI:** 10.1101/2020.01.22.914051

**Authors:** Nasim Bararpour, Federica Gilardi, Cristian Carmeli, Jonathan Sidibe, Julijana Ivanisevic, Tiziana Caputo, Marc Augsburger, Silke Grabherr, Béatrice Desvergne, Nicolas Guex, Murielle Bochud, Aurelien Thomas

## Abstract

As a powerful phenotyping technology, metabolomics provides new opportunities in biomarker discovery through metabolome-wide association studies (MWAS) and identification of metabolites having regulatory effect in various biological processes. While MS-based metabolomics assays are endowed with high-throughput and sensitivity, large-scale MWAS are doomed to long-term data acquisition generating an overtime-analytical signal drift that can hinder the uncovering of true biologically relevant changes.

We developed “*dbnorm*”, a package in R environment, which allows visualization and removal of signal heterogeneity from large metabolomics datasets. “*dbnorm*” integrates advanced statistical tools to inspect dataset structure, at both macroscopic (sample batch) and microscopic (metabolic features) scales. To compare model performance on data correction, “*dbnorm*” assigns a score, which allows the straightforward identification of the best fitting model for each dataset. Herein, we show how “*dbnorm*” efficiently removes signal drift among batches to capture the true biological heterogeneity of data in two large-scale metabolomics studies.

## Introduction

Large metabolome datasets, generated by metabolomics assays, have become an essential source of information about molecular phenotypes in system biology studies ^1-3^. As an intermediate molecular layer between genes and disease phenotypes, metabolite levels and the corresponding pattern of changes are highly associated with the degree of perturbation in biological systems and rewired metabolic networks in a given phenotype ^41-12^.

Liquid chromatography-MS (LC-MS) has become the most widely used experimental platform in the study of the metabolome, owing to the progressive improvement of the instrumental conditions in terms of sensitivity and selectivity ^13^. However, LC-MS based metabolomics assays suffer from the inherent variation in the distribution of signal measurements and/or in the signal sensitivity and intensity driven by external factors^13^. This signal drift remains a major limitation to data normalization in biomedical and clinical studies, adding up to the biological inter-individual variability and unavoidable technical variation introduced during sample preparation^14^. In particular, such drift can significantly compromise the technical precision and signal stability in large-scale studies, where the data acquisition for several hundred to thousands of biological samples needs to be done in different analytical blocks (i.e. batches) over several weeks or even months^14,15^. In this case the largest variance in the dataset may be assigned to the batch effect or experimental run order, thus hindering the real biological difference and true functional signals, leading to data misinterpretation ^14,16,17^. Therefore, prior to any chemometrics analysis, large metabolomic datasets need to be corrected for the unwanted analytical, within- and between-batch variation, in order to make data comparable and reveal biologically relevant changes ^18^.

In most of the LC-MS-based metabolomics data processing workflows, signal intensity drift correction is performed using quality control (QC) samples per metabolite feature. QC samples are aliquots of a QC pool, representative of entire sample set. QC samples are injected within each batch periodically (every 4-10 samples) to monitor signal drift over time and evaluate the system performance and data quality^19-21^. However, in large-scale studies, the preparation of such QC samples may be difficult due to large number of samples, whose handling would involve additional freeze-thaw cycles. Moreover, one may want to start the data acquisition before finalizing the sample collection. In such cases, surrogate QC samples are required ^19,22-25^. In general, the drift correction using QCs is based on the assumption that the same sources of variation apply to the metabolites present in both biological samples and QCs, as their representative pools. In a majority of QC-based workflow, an equalization step is considered for batch-effect correction to remove signal intensity drift via several commonly proposed algorithms, such as batch-ratio-based correction ^26^, regression-based models ^19,26,27^ and/or linear and non-linear smoothing algorithms (e.g. *lowess*-model)^19,27-30^. Beyond the algorithm-specific assumptions, the fundamental premise of all QC-based correction models is that QCs and study samples contain the same metabolite features ^23,27^, a condition that might not be met when using surrogate QC sample. In this case, the features that are detected in the subject of study, but are missing in the QCs, must be excluded before the application of a QC-based correction model. This limitation leads to loss of information in the metabolic phenotyping and may bias new discoveries ^19,31,32^.

Another implemented solution for correcting signal drift in the analytical measurements via LC-MS-based metabolomics is spiking stable isotope-labeled metabolites into samples prior to metabolite extraction. This procedure is supposed to control both signal fluctuation during cross-comparison of different batches and biases introduced during sample preparation ^33,34^. However, metabolome size and diversity are too high to be completely covered by corresponding internal standards^35^, and signal variability for internal control metabolites may not truly reflect that of other endogenous or /and exogenous compounds, due to the difference in their chemical properties ^25,36^. Hence, this method should not be favored for the correction of potential batch effects in metabolomics assays ^37^.

Alternatively, using models and algorithms that are not dependent on QCs might be a genuine and alternative way to compensate for technical sources of variation in a large-scale/long-run study. In particular, there has been a growing interest in adopting and applying specific statistical methods that were originally developed for microarray based gene expression data to adjust for unwanted variation in metabolomics data. Such methods include removal of unwanted variation (*RUV*) model^38-40^, linear model for microarray data (*LIMMA*)^36,41^, and *ComBat* using empirical Bayes methods (*EB*) ^17,42-44,45^. While these models can successfully correct for unwanted variation, they rely on a set of assumptions limiting their application to metabolomics data. For example, on one hand *RUV* model depends on prior knowledge of metabolome features (variables) detected in the study groups, because it settles the analytical variation based on the behavior of negative control features, whose levels are supposed to remain constant in different biological conditions ^36,39^. On the other hand, *LIMMA* function strictly dependent on some additional information on sample metadata such as biological covariates, which are used for the design of the model matrix ^41^. In contrast, *ComBat* has the flexibility to adjust batch effect by integrating information on both batches and biological covariates or just on batches if demography of population is not available ^9,14^. This function is developed in the global term of “Parametric shrinkage adjustment” and accommodates batch effect in three steps: data standardization, batch effect estimation via empirical priors and batch effect removal using the adjusted estimators ^45^. Although making strong parametric estimation, *ComBat* has been recognized for its superior performance in adjustment of unwanted variation compared to several others models ^42^. *ber* is another statistical model that was developed for batch effect correction in gene expression data. It is based on a two-stage regression procedure and uses linear fitting for both location and scale (L/S) parameters ^46^. Interestingly, this model showed a better performance in microarray data compared to the empirical Bayes model implemented in *ber*, but was never tested for metabolomics data ^46^.

Herein, we describe “*dbnorm*”, a new package in R environment, which incorporates a collection of functions applicable to large-scale metabolomics datasets for data visualization and for normalization cross batches.

## Results

### 1. Statistical modeling for intra- and inter-batch signal drift correction in large-scale metabolomics datasets

Despite their efficiency in removing intra- and inter-batch signal drift, the application of QC-based approaches might be limited by the impossibility to have QC samples highly representative of the whole sample set, particularly in large-scale population studies. To overcome this issue, statistical models that were originally developed for microarray gene expression data can be used to address the unwanted variation (e.g. batch effect) in metabolomics data. These statistical models offer the flexibility to accommodate signal variation across multiple batches based only on the sequence of acquisition, or integrating also other available population demographic characteristics (i.e. treatment)^9,14^, which is recommended in case of uneven study design.

We developed a new R package “*dbnorm*” in which, besides several functions for pre-processing of data and estimation of missing values, we assembled two distinct functions for batch effect correction based on statistical models: *ComBat* (both *parametric* and *non-parametric*), that was already in use for metabolomics data normalization, and *ber*, that we propose here as an additional tool for drift adjustment across multiple batches in metabolomics datasets. The performance of these two statistical models was assessed on two different big metabolomics datasets, using traditional QC-based methods (ie. *lowess -*model) as a reference for result comparison. Of note, for each analyzed dataset, “*dbnorm*” generates diagnosis plots and calculates a score value which helps users to easily choose the statistical correction model which best fits their data structure. The statistical models and the underlying algorithms are explained in the method section. The implementation of this package is publicly available at https://github.com/NBDZ/dbnorm

### 2. Drift correction in a large-scale targeted metabolomics dataset from a human prospective cohort study

The first dataset employed to test “*dbnorm*” is a set of targeted metabolites from plasma samples of 1,079 individuals, analyzed in 11 analytical batches over a period of 12 months, and yielding data on 239 metabolites detected across all samples (see method section).

#### 2.1 Across-batch drift assessment

As previously described, in the majority of large-scale metabolomics experiments, QCs are periodically analyzed throughout an analytical run to allow for signal drift and data quality assessment. Therefore, distribution of QC signal is indicative of amplitude of analytical variation in the analyzed dataset. To perform the QC-based drift correction on the above specified dataset acquired on human cohort, we retrieved the data for 135 QCs injected periodically, every 10 samples. Unsupervised principal component analysis (PCA) of acquired individual metabolic signatures, including 239 detected metabolites, revealed a clear distinction between QC sets analyzed in different batches or experimental runs (Fig.1A), with the first two principal components (PCs) explaining 65% of the variance mainly derived from batch effect or analytical variability (Supplementary Figure 1A).

**Figure 1.**
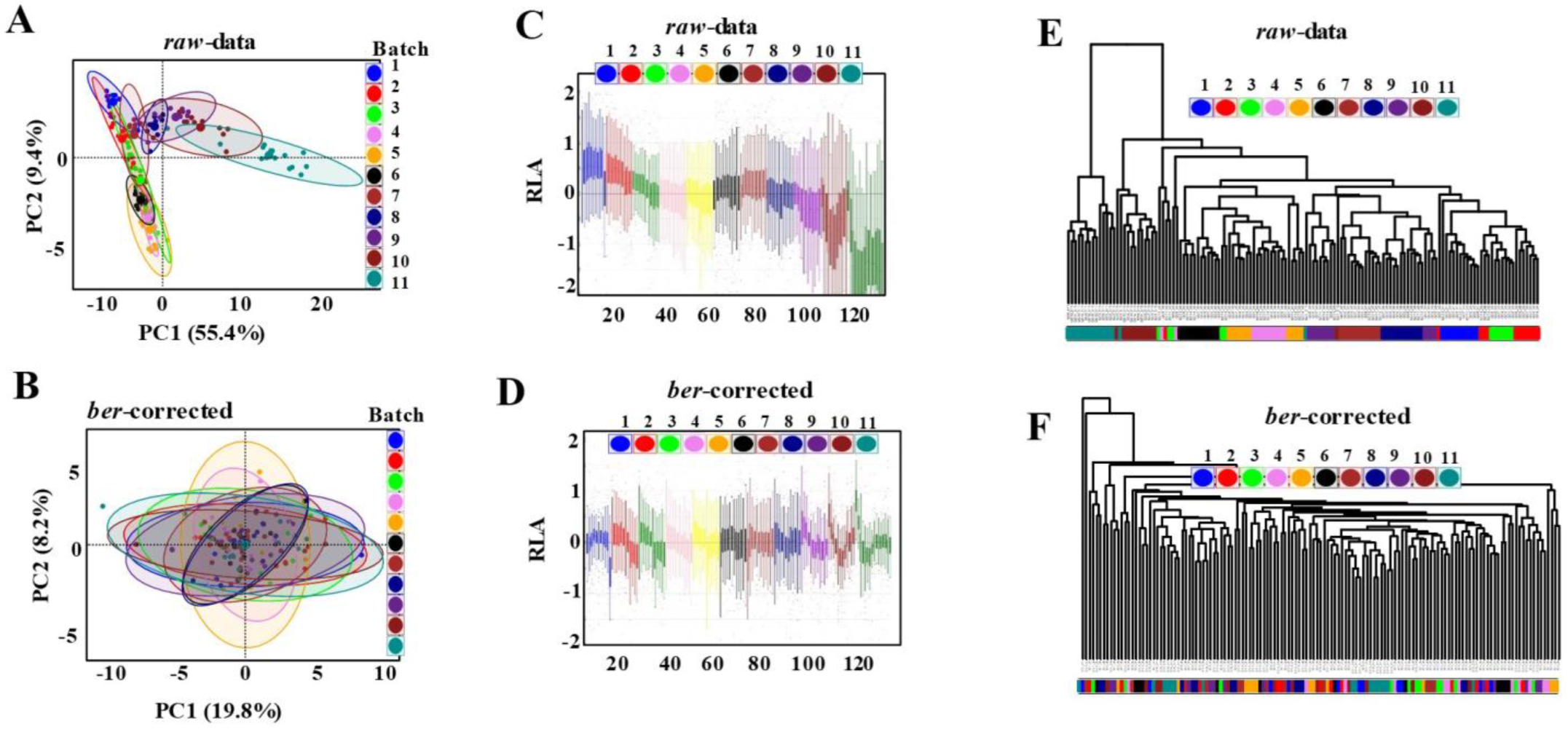
Detection of batch effect in LC-MS targeted metabolomics analysis of QC samples of a human cross-sectional study. Principal components analysis (PCA) of plasma metabolome in QC samples shows that in raw data the main source of variance is the inter-batch variability **(A)**, which is removed after adjustment through *ber*- model **(B)**. Relative log abundance (RLA) plots of data showed average distribution of the metabolites in QC samples before **(C)** and after batch effect removal using ber-model **(D)**. For corrected data RLA median is close to zero and shows lower variation compared to raw data. Dendrogram of raw data **(E)** clearly visualizes the batch-dependent pattern on QC samples (clustered sets of samples according to the analytical run), while this pattern is abolished in *ber*-corrected data **(F)**. The colors are indicative of the batch number. Graphs are generated on LC-MS data obtained in positive polarity mode along 11 batches of analysis.

Importantly, total ion current (TIC) in the QCs showed a 10-fold signal intensity drift across batches, with a gradual signal decrease from 1.3 10^7^ ion counts in batch 1 to 5 10^6^ ion counts in batch 11 (Supplementary Figure 2). Consistently with that, the relative log abundance (RLA) plot showed a moderate drift in the total centered median ion intensity across the experimental runs (Figure 1C). Moreover, multivariate unsupervised hierarchical clustering analysis (HCA) confirmed the sample clustering per batch, as depicted in Figure 1E. To correct for within and between batch effects and normalize the data prior to statistical analysis, we applied the statistical models implemented in “*dbnorm*” R package, namely *ber* and *ComBat* (both parametric and non-parametric versions). PCA was applied to assess the efficiency of drift correction. The score plot showing the centered cluster of QCs between different batches was indicative of batch effect removal, with variance explained by PC1 and 2 reduced to < 40 % (Figure 1B and Supplementary Figure 1B, 3A, B). In parallel, across-batch RLA plot showed the adjusted median signal intensity between batches (Figure 1D, 1F).

Following the data quality assessment using QCs, we performed explorative statistical analysis of biological samples comprising the analyzed individuals, prior to and following the drift correction. PCA plot of raw data (i.e. prior to drift correction) generated in both positive and negative ionization modes showed sample clustering by batch order (Figure 2A). In addition, probability density function plots (PDF plots) of metabolite distribution across batches showed shifted PDFs for the majority of metabolites detected in the study samples (see example of 3-Hydroxy-3-methylglutarate and 2’,3’-cyclic phosphate 3’-CMP in Supplementary Figure 4A and B). In parallel, Adjusted-Coefficient of Determination (Adj. R. squared) revealed the high dependency between variability in the dataset and across-batch signal drift, with > 50% of variability explained by batch for most metabolites (Figure 2D and Supplementary Table 1).

**Figure 2.**
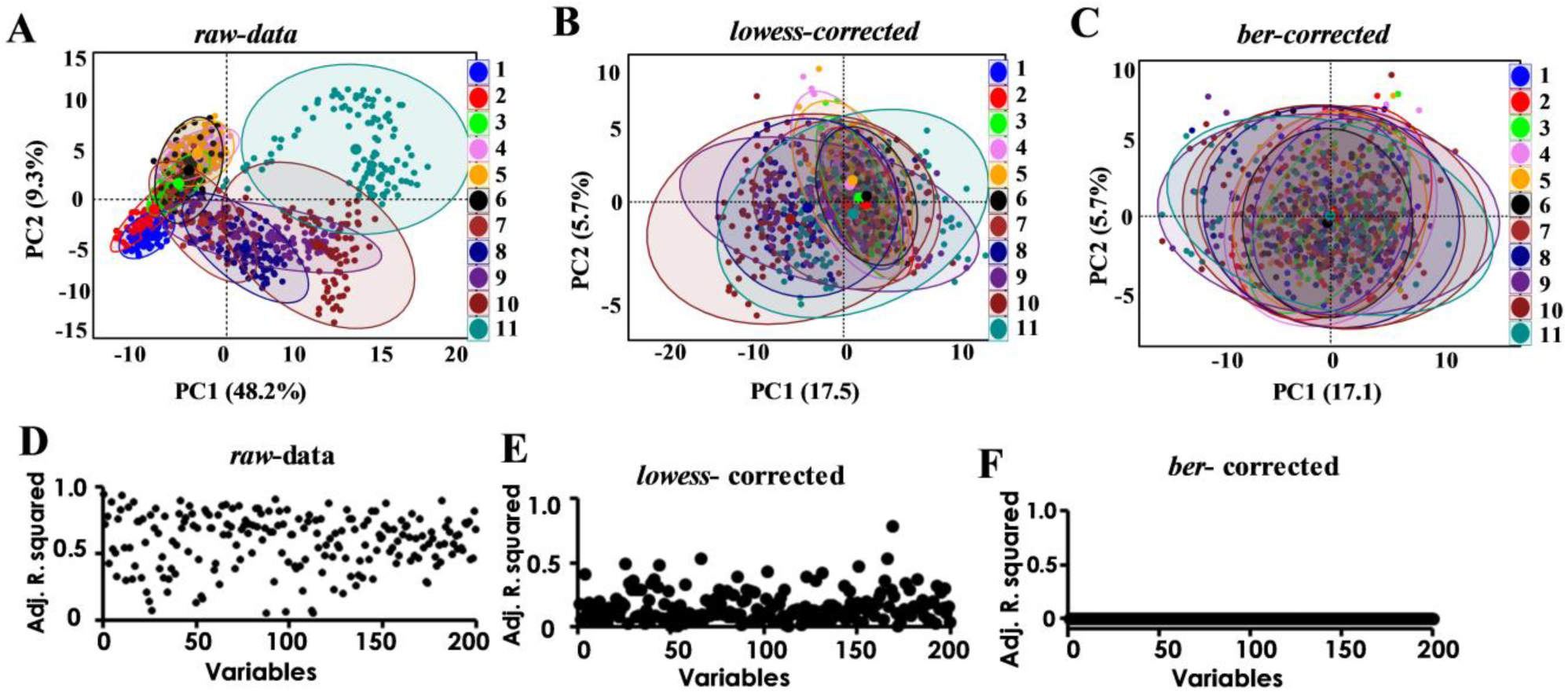
Detection of batch effect and its correction in LC-MS targeted metabolomics analysis of 1079 samples of a human cross-sectional study. 1079 plasma samples were analyzed by LC-MS targeted metabolomics in 11 analytical batches. Principal components analysis (PCA) of raw data **(A)** shows separation of sample clusters mainly triggered by batch order. This separation is removed after adjustment through *lowess* **(B)** or *ber*- model **(C)**. Adjusted Coefficient of Determination (Adjusted R-squared) estimated by regression model shows the dependency of dataset variability to batch level. The raw dataset presents high variability **(D**), which is completely removed in the dataset corrected by using *ber* -statistical model (F), while *lowess*-corrected data **(E)** still shows some level of association between batch and some metabolites, with the highest dependency to the level of 78%.

We thus employed the three available statistical models implemented in the “*dbnorm*” R package (*ber, parametric* and *non-parametric ComBat*) to accommodate the signal drift observed in the whole dataset. To evaluate their efficiency to correct the signal drift within- and between-batches, we compared them to reference QC-based models, such as *lowess*. In general, the spatial separation predictive of batch effect was significantly reduced after correction of data with all tested models (Figures 2B-C and Supplementary Figure 3C-D). By applying both types of correction algorithms (QC–based and non-QC based), the largest variance captured by PC1 decreased from about 50% observed in the raw dataset to less than 20 %. Besides, overlapped PDFs plots was observed after data correction using either QC-based model (Supplementary Figure 4 C,D) or statistical models (Supplementary Figure 4 E-J). Of note, *lowess*-corrected dataset lacked xanthosine 5’-monophosphate metabolite when compared to the datasets generated by statistical models. This metabolite is a good example of a low abundant metabolite, likely present in only one portion of samples and thus below the limit of detection in QC samples due to the pooling - dilution effect.

Although the QC-based *lowess* model considerably reduced the batch-associated signal variability, as indicated by the decrease of the Adj. R. squared value compared to raw data (Figure 2E), the signal intensity for some metabolites was still strongly associated with the injection order over time estimated by regression model (Figure 2E, Supplementary Table 2). As an example, the clear batch-dependent shift in the signal intensity of citrate that was observed in the raw data, was preserved for some batches in the *lowess*-corrected data (Supplementary Figure 5A, B). In contrast, in all datasets that were corrected with one of the “*dbnorm*” statistical methods (*ber, parametric* and *non-parametric* models), the average intensity of this metabolite remained constant across batches (Supplementary Figure 5C-E). Accordingly, the use of statistical models improved to a greater extent the decrease in the batch-associated signal intensity drift compared to QC-based model (Figure 2F, Supplementary Figure 6 and Supplementary Table 3,4,5).

Depending on proportion of variability explained by the batch, the “*dbnorm*” tool will calculate a score for the maximum variability defined in each model to facilitate the conclusion about the model which provides the best compromise for drift correction considering the consistency of overall model performance for all detected metabolites. As shown in Supplementary Figure 7, in this cohort study, the maximum variability detected for a metabolite was at 0.78 (78%) for the *lowess*-corrected dataset. Similarly, for the *non-parametric ComBat* corrected dataset, a residual maximum variation of 0.60 (60%) was still detected (Supplementary Table 5 indicating the remaining, for some metabolites, of the signal drift across batches. In contrast, the datasets corrected by *ber*- and *parametric ComBat* -models presented a similar performance in this study, with a maximum variability associated to the batch < 0.01 (1%) (Supplementary Table 3,4). These results indicates that *ber*- and parametric models are more efficient in normalizing the signal intensity changes in this large-scale metabolomics dataset.

#### 2.2 Treatment of batch effects improves downstream differential analysis

To investigate to which extent the quality of data might affect the data interpretation in this study, we next explored the impact of each type of correction algorithms on the downstream statistical and metabolic pathway analysis.

All the subjects of this prospective cohort study were previously phenotyped for kidney functionality and associated parameters, such as age, sex, and creatinine clearance, as an indicator of renal impairment to estimate the severity of a kidney disease^47^. In addition, Glomerular Filtration Rate (GFR), a major surrogate of kidney function, is measured on a continuous scale and estimated by Chronic Kidney Disease Epidemiology Collaboration (CKD-epi)^48,49^ (Methods). We thus took advantage of this information to have further indication on the performance of the different statistical models employed for data correction. We first evaluated the correlation between creatinine levels, determined by the targeted LC-MS/MS metabolomics experiment with that of measured at clinical laboratory based on an enzymatic assay. We looked at 5 datasets: raw data, *lowess-* corrected, *ber-*corrected and *parametric*- and *nonparametric*-*ComBat* -corrected. All applied normalization models improved the correlation between creatinine levels measured clinically and those obtained using targeted LC-MS/MS analysis. The correlation coefficient (Pearson’s r) passed from 0.18 (Fisher’s z= 0.18 with 95% interval level (**CI**), 0.12 to 0.24) observed in raw MS data, to 0.5 (Fisher’s z= 0.54 with 95% CI, 0.48 to 0.60) in the *lowess*-corrected data and up to 0.61 (Fisher’s z= 0.71 with 95% CI, 0.64 to 0.76) for the data corrected by statistical models (Figure 3A). Using a multiple regression model, we found that the expected significant correlation between the outcome of renal failure such as CKD-epi and creatinine (predictor) can be revealed only when data are corrected for batch effect through either QC-based or statistical-based models, as shown by the increased absolute regression coefficients and decreased p-value in both *lowess*-and *ber*-corrected data (Figure 3B). Likewise, other metabolites that are known to be associated to renal failure such as gluconate, citrulline, and hippurate^47^, showed a significant association to CKD-epi level only in the *lowess-* or *ber-* adjusted data (data not shown).

**Figure 3.**
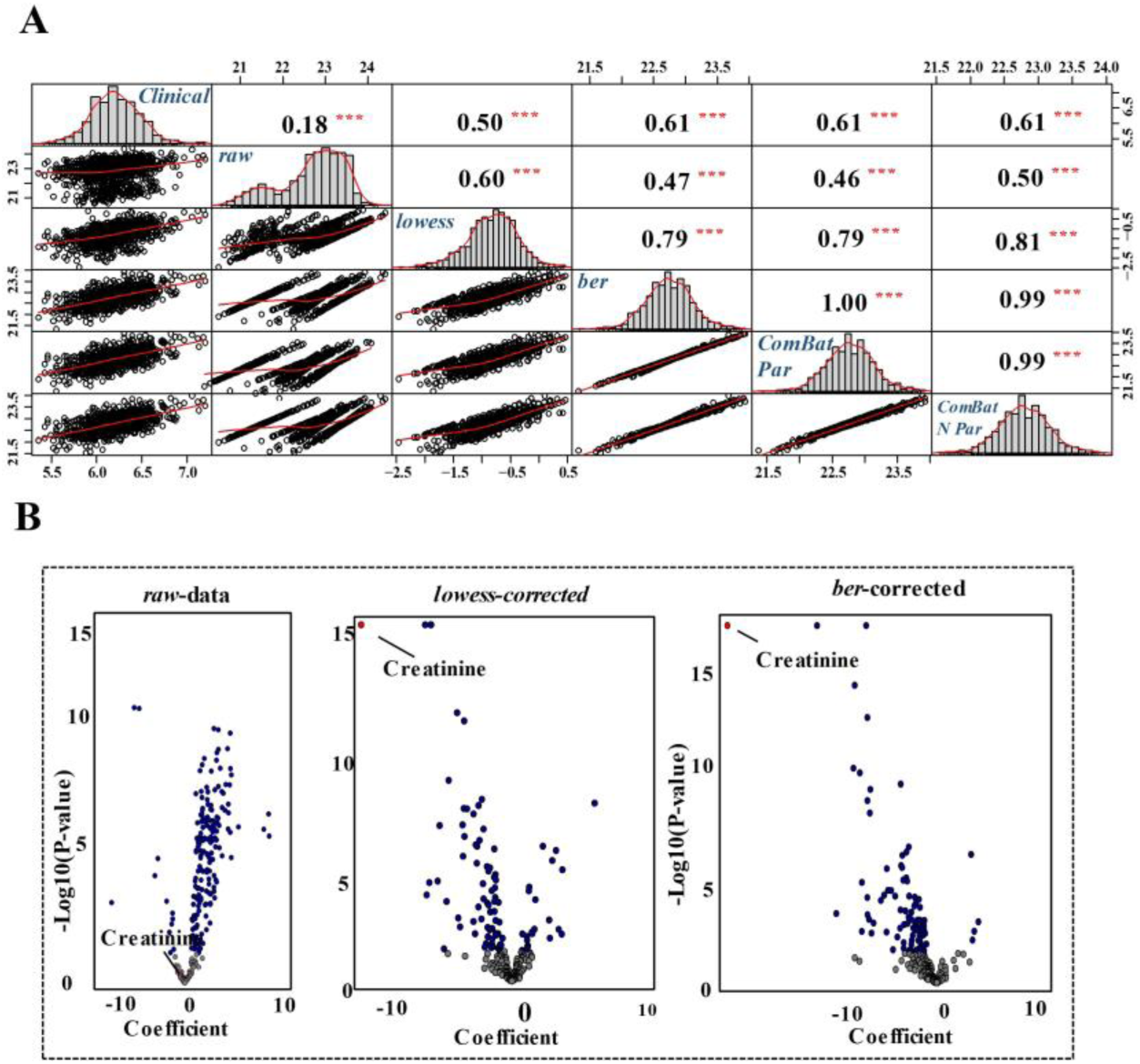
Impact of the correction of batch effect in LC-MS targeted metabolomics analysis of a human cross-sectional study on downstream functional analyses. Pearson correlation between levels of creatinine as measured by clinical test and its levels, as measured in LC-MS metabolomics analysis, before or after batch effect correction **(A)**. The correlation (Pearson’s r) increases from 0.18 in the raw dataset, to 0.5 in the *lowess*-corrected dataset, to a maximum level of 0.61 in the datasets corrected via statistical models (*ber, ComBat* – par - and *ComBat*-*N par*). *** indicates p-value <0.001. **(B)** Association of CKD-epi and metabolite levels fails in detecting significant association with creatinine levels in raw data, while this expected association is observed in both lowess- and ber-corrected datasets. The model is adjusted by age and sex.

### 3. Drift correction in a large-scale untargeted metabolomics dataset from a mouse model study

#### 3.1 Across batch drift assessment

Untargeted metabolome profiling is mostly employed to increase the chance of identifying unexpected discriminant biological signal, as it allows for the detection of as many metabolite features as possible from diverse chemical classes, without an a priori hypothesis. Due to high levels of noise and redundancy, the use of a robust statistical model for batch effect correction is even more important in untargeted metabolomics datasets. Here, the aim was to evaluate further the efficiency of *ber*-, *parametric*- and *non*-*parametric ComBat* models, implemented in “*dbnorm*” tool, for batch effect removal in a dataset acquired in a full scan mode in an untargeted assay.

Briefly, the metabolome profile of two types of adipose tissue, visceral (v-AT) and subcutaneous (sc-AT), was measured in mice fed with a high fat diet (HFD) and/or control diet (ctrl). The diet treatment was scheduled for 1 and 8 weeks. Overall, 264 samples, including QCs, were analyzed in a full scan mode in an untargeted metabolomic assay with the data acquisition divided in three analytical batches. Data processing using XCMS software (https://xcmsonline.scripps.edu/) yielded 11,156 aligned metabolite features defining multiparametric metabolic signatures (or profiles). Unsupervised multivariate analysis of these metabolic profiles (i.e. PCA and HCA) supported the presence of a strong batch effect, with samples clustered in the corresponding analytical batches (Figure 4A). In addition, the adjusted coefficient of variation estimated by regression model indicated that the variability of certain metabolites could be entirely explained by the signal intensity drift between batches (Supplementary Figure 8B). Following the signal drift correction using statistical models, the batch effect was removed, as suggested by the PCA scores plot and HCA dendrogram (Figure 4B and D, Supplementary Figure 8A). The high variability observed in the raw dataset was strongly reduced after batch effect correction via statistical models for the majority of variables (metabolites), although a maximum variability of about 20% was still present for some variables in the data corrected by non-parametric *ComBat* model (Supplementary Figure 8E, F). Interestingly, as compared to parametric *ComBat*-corrected represented in auto-scaled graph (Supplementary Figure 8D, F), the *ber*-corrected dataset showed a lower dependency to the batch order, with a consistent effect on all variables (metabolites), as demonstrated by the Adjusted R-squared which was almost zero for all the detected features (Supplementary Figure 8C, F). The very low negative Adjusted R-squared values detected here are usually indicative of very poor fitted regression model, indicating a weak dependence to the signal drift or batch effect estimated by the model in the *ber*-dataset.

**Figure 4.**
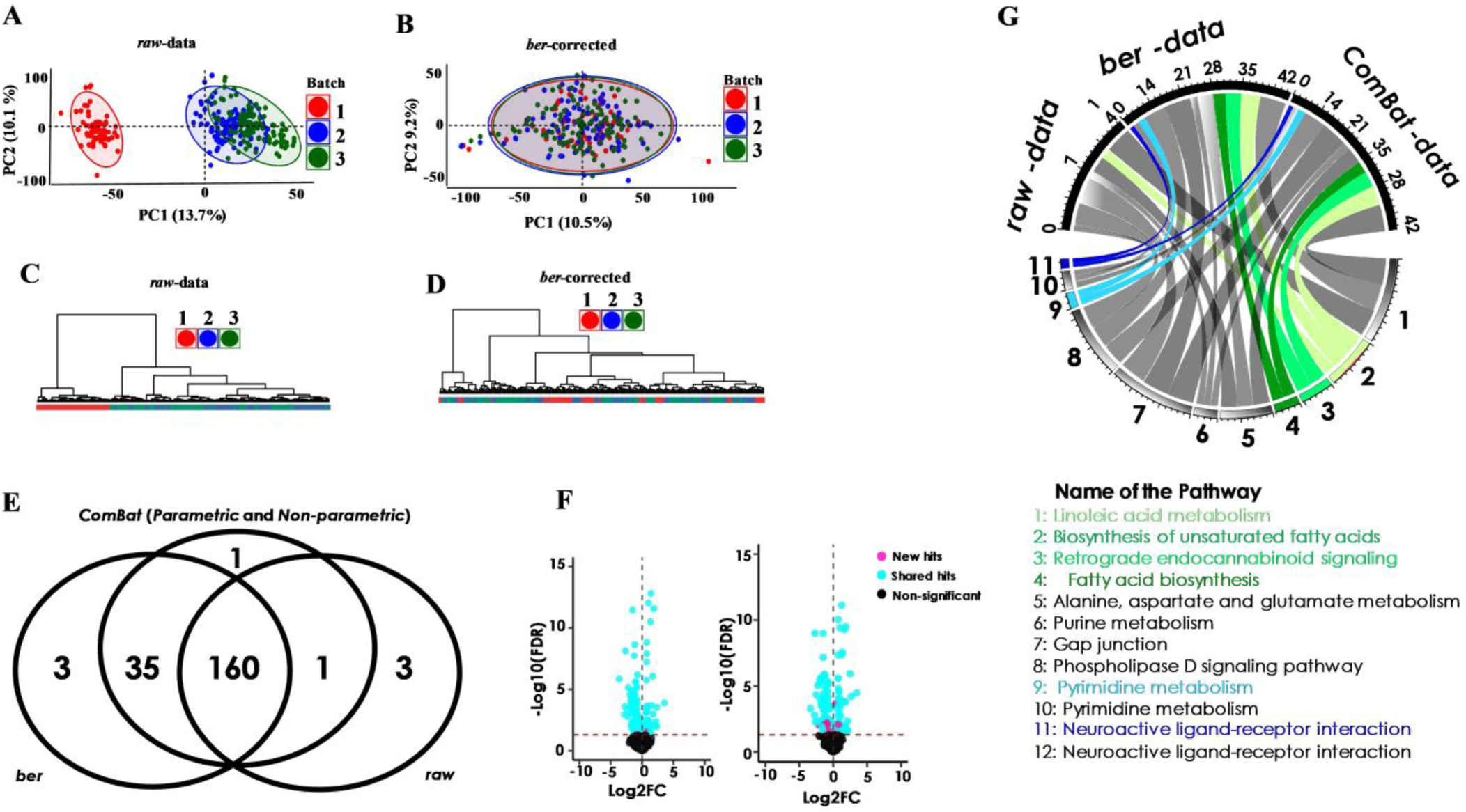
Detection of batch effect and its correction in LC-MS untargeted metabolomics analysis of 264 adipose tissue samples and its impact on downstream analyses. PCA plot of mouse fat tissue metabolome acquired in negative polarity shows separation among the samples associated with the experimental run in raw data **(A)** which is removed after adjusting for batch effect using *ber* statistical model **(B)**. Dendrogram of raw data shows sample clusters particularly for batch 1 **(C)**, which are not seen anymore after batch effect removal using *ber*-function **(D)**. The color core indicates the batch number. **(E)** Venn diagram shows the number of metabolites significantly altered by 8 weeks of HFD (with FDR < 0.05) before and after batch correction through statistical models (*ber, ComBat-par* and ComBat-*Npar*). Correction with either *parametric* or *non-parametric ComBat* results in the same list of differentially expressed metabolites. **(F)** Volcano plots of raw and *ber*-corrected data show the impact of HFD on the list of differentially expressed metabolites. The metabolites appearing as significantly regulated only in *ber*-corrected data are depicted as pink dots. **(G)** Circos plot showing the relative size of the metabolic pathways that are enriched in the different lists of metabolites differentially regulated in response to HFD, obtained before (raw data) or after correction for batch effect using either *ber* or *ComBat* statistical models. The circle’s segment on the top indicates the dataset type, with the differentially regulated metabolites defined in each dataset. Numbers on the bottom of the circle indicate the metabolic pathway, as defined in the legend below. The connecting bands link the enriched biological functions to the underlying differentially regulated metabolites. Grey connecting bands are associated to biological pathways enriched in all datasets. Green and blue connecting bands highlight biological pathways that are enriched only in *ber-* or *ComBat*-corrected datasets.

#### 3.2 Batch effect correction facilitates the discovery of biologically relevant changes

We finally investigated the importance of data correction for unwanted variation in a large-scale untargeted metabolomics assay with the aim of capturing true biological differences due to treatment in our case study. To this end, we compared the candidate list obtained from statistical analysis of raw data to that resulting from normalized data following the batch correction using the statistical models *ber, parametric* and *non-parametric ComBat*, implemented in the “*dbnorm*” tool. While the majority of the significant changes were detected in all analyzed datasets, the list of candidate metabolites associated with the HFD treatment was more exhaustive after adjustment (Figure 4E). As depicted in the volcano plot (Figure 4F), the levels of several metabolite features (variables) were revealed as significantly different only after batch effect removal, thus suggesting that in the raw dataset some biological signals induced by HFD might be overlooked due to high analytical variability. Accordingly, the data interpretation using metabolite set enrichment analysis (MSEA) varied between raw and normalized datasets. For instance, several biological pathways including “Fatty acid biosynthesis”, “Retrograde endocannabinoid signaling” and “Biosynthesis of unsaturated fatty acids”, “Linoleic acid metabolism “, that were previously shown to be influenced by HFD in adipose tissue ^50-52^, were enriched only using the list of candidates as a result of statistical analysis of normalized datasets (Figure 4G). No significant difference was observed depending on the statistical model applied thus cross-validating the performance in drift correction.

## Discussion

Metabolic profiling offers holistic determination of intermediate and end products of metabolism whose deviation from normal level might provide important information on the dysregulation of metabolic pathways in disease condition. Such changes can be caused by genetic disorders, environmental factors, drug treatment, etc. This information could help improving disease diagnosis, prognosis and treatment choice. However, population-based studies need great care of experimental design and post hoc equalization model to generate comparable sample sets across batches. Periodical injection of QC samples is one of the most commonly used methodology in the metabolomics community and is exploited by QC-based normalization models to correct for drift across batch effects. However, this methodology has the limitation of being dependent on the availability of QCs truly representative of all the study samples, whose adequate preparation is not always possible, particularly in large-scale studies. If surrogate QC-samples are used, QC-based normalization methods might fail in removing batch effect homogenously for all the features characterizing the multiparametric metabolic profile. Our results confirm the limits of such correction methods. For instance, we show that in the human dataset that we analyze, citrate levels remain highly associated to the batch order in the *lowess*-adjusted data, which might generate bias in data interpretation. On top of that, the analysis with a QC-based correction model is restricted only to the metabolites that are detected in both QC and study samples, thus potentially impairing the discovery of novel biological hits in a medical condition. In our dataset, xanthosine 5’-monophosphate is only detected in the study samples, but not in QCs, and is therefore discarded a priori by the *lowess*-model.

In our human population dataset, while variability linked to batch is still present for some metabolites by using QC-based *lowess* correction model, all the statistical models implemented in the “*dbnorm*” package, present a higher performance on the overall correction of signal drift across batches, with *parametric ComBat* and *ber* showing the best score. However, we cannot exclude that other datasets, with a different data structure, might be more effectively adjusted by *non parametric ComBat* and QC-based models.

Our results clearly demonstrate the substantive impact of data adjustment for analytical heterogeneity on the prediction of clinical outcomes. In the human study, data normalization triggered an increased association between eGFR (the outcome measured via the CKD-epi formula) and creatinine, thus highlighting a pattern that was not detected in raw data. This also indicates that data correction performed on our dataset is not overfitting the data, but rather favoring the detection of biologically relevant differences. Knowing that *ComBat*-model is developed to avoid overfitting of the data in case of small sample set in each batch, we found computational advantages using the *ber*-model, thanks to higher speed in processing the metabolomic data.

Similarly, in our mouse experiment, the statistical models compensating for across batch signal drift drastically decreased the high variability associated to batch level in the raw dataset to the almost zero level in the corrected datasets, with a more consistent removal observed when employing *parametric ComBat*-model and *ber*-model. Data correction and automation resulted in a slightly distinct list of differential features associated with HFD-treatment, with similar candidates given by *ber*-, *parametric*-*ComBat* and *non-parametric*-*ComBat* models. Although the list generated by using raw data lacked only few metabolite candidates, this small difference impaired the enrichment of a series of biological pathways that are known to be affected by HFD treatment ^46,48,49^, thus reducing the biological significance of the results.

In conclusion, in agreement with previous reports, our study supports the necessity of data cleaning from unwanted technical variation, which helps improving the detection of biological mechanisms underlying a treatment or medical state. “*dbnorm*” is an efficient and user-friendly tool for removal of drift across batches. It helps users to diagnose the presence of analytical drift thanks to several visual inspections based on advanced statistical inferences implemented in the package. In addition, different statistical models are implemented, namely *ComBat*- and *ber*-. In addition, several functions implemented in the “*dbnorm*” assist users to visualize the structure of large datasets after correction via the implemented methods, distinctively from the perspective of samples analyzed in the entire experiments and the features detected in the study samples. Notably, “*dbnorm*” and its application is not limited to the metabolomics data, but could be extended to other high-throughput techniques.

## Methods

### Package

“*dbnorm*” and its functions are explained in details in the Package Manual. Briefly, it includes distinct functions for pre-processing of data and estimation of missing values, conventional functions for batch effect correction based on statistical models, as well as functions using advanced statistical tools to generate several diagnosis plots to inform users about their data structure. The “*dbnorm*” package includes statistical tools which allows user to inspect the structure and quality of multidimensional datasets of large metabolomics datasets at both macroscopic and microscopic scale, namely at the sample batch level and metabolic feature level, respectively.

Batch correction models implemented in the “*dbnorm*” are adapted from microarray analysis, namely, *ber*-model, a package from CRAN (ber: https://cran.r-project.org) and *ComBat*-model with both parametric and non-parametric setting, from sva package in R (sva; https://bioconductor.org). In brief, *ComBat* uses EB method to remove location (mean) and scale (variance) of batch effect. Notably, EB model borrows information across genes and experimental conditions and assumes that systematic biases (i.e. batch effect) often influences many genes in the same way. Then it estimates variance for each gene within batch and across batches. And finally, standardized data (*Z*_*ijg*_) is calculated via

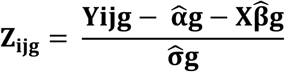

In which *Yijg* is expression value for gene *g* for sample *j* from batch *i, 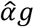* is the overall gene expression, *X* is a design matrix for sample conditions, and 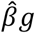 is the vector of regression coefficients corresponding to *X* and estimated variance of 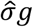.

In contrast, *ber*-function uses linear regression at two stages to estimate location and /or scales parameters:

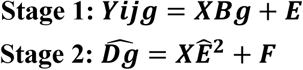

In which *Bb* which is *mb* × *g* matrix for batch *l*= 1,…,*mb* and gene in *j* = 1,…,*g* is estimated by first regression model where E is a matrix of error. The second regression is applied on the squared residual of the first stage, *Ê*^2^. 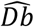 is the estimated matrix parameter for the scale batch. Upon calculation of its mean denoted by 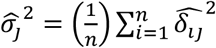 for gene *j*, data are transformed to calculate and remove batch effect. This package is publicly available at https://github.com/NBDZ/dbnorm.

### Human study

SKIPOGH (Swiss Kidney Project on Genes in Hypertension) [grant number: 33CM30-124087] is a family-based multi-center population-based study exploring the role of genes and kidney haemodynamics in blood pressure (BP) regulation and kidney function. Method and population are described in details elsewhere ^53-55^.

### Animal experimentation

All animal experiments [grant number: 310030_156771] and procedures were approved by the Swiss Veterinary Office (VD-2942.b. and VD-3378). C57/BL6 male mice were purchased from Janvier Labs and housed 5 per cage in the animal facility of Centre for Integrative Genomics, University of Lausanne.

Four-week old mice were fed for two weeks with a 10% in fat chow diet (D12450J, Research Diet). At 6 weeks of age they were either shifted to a high-fat diet (HFD) containing 60% fat (D12492, Research Diet) or kept on a control diet for 1 or 8 weeks. Random blocking was used. Efficiency of the diet-induced obesity was followed by regular measurements of weight^56^. All animals were kept in a 12:12 h light:dark cycle with water and food ad libitum. All the mice were sacrificed by CO2 between ZT2 and ZT5. In this study sc-AT refers to inguinal subcutaneous adipose tissue in mice.

### Metabolomics

Targeted metabolomics analysis conducted on the plasma samples of the human prospective cohort study study. Metabolites were extracted from 100 µL plasma samples using a methanol-ethanol solvent mixture in a 1:1 ratio. After protein precipitation, supernatant was evaporated to dryness and finally re-suspended to 100µL H2O 10% MeOH. The samples were analyzed by LC-MRM/MS on a hybrid triple quadrupole-linear ion trap QqQ_LIT_ (Qtrap 5500, Sciex) hyphenated to a LC Dionex Ultimate 3000 (Dionex, Thermo Scientific). Analysis were performed in positive and negative electrospray ionization using a TurboV ion source. The MRM/MS method included 299 and 284 transitions in positive and negative mode respectively, corresponding to 583 endogenous metabolites. The Mass Spectrometry Metabolite Library (Sigma Aldrich) was used as reference material for the standard metabolites.

The chromatographic separation was performed on a column Kinetex C18 (100×2.1 mm, 2.6 µm). The mobile phases were constituted by A: H2O with 0.1% FA and B: ACN with 0.1% FA for the positive mode. In the negative mode, the mobile phases were constituted by A: ammonium fluoride 0.5 mM in H2O and B: ammonium fluoride 0.5 mM in ACN.

The linear gradient program was 0-1.5 min 2%B, 1.5-15 min up to 98%B, 15-17 min held at 98% B, 17.5 min down to 2%B at a flow rate of 250 µL/min.

Total 1081 different plasma samples were analyzed in 11 batches over 12 months. 135 and 156 QCs were periodically injected between sample runs in the positive- and negative-modes respectively. Surrogated QCs were considered in this study to prevent repeated thawing-refreezing cycles.

The MS instrument was controlled by Analyst software v.1.6.2 (AB Sciex). Peak integration was performed with MultiQuant software v.3.0 (AB Sciex). The integration algorithm was MQ4 with a Gaussian smoothing of a half-width equal to 1.5 points.

To obtain the fat metabolome profile in the mice, metabolites were extracted from 10-20 mg of fat depots either sc-AT or v-AT using 400 µL of mix organic solvent comprising EtOH: MeOH: H2O in the proportion of 2:2:1 to remove protein efficiently as well as to extract polar and semi-polar metabolites successfully. All the samples were then vortexed mixed for 30 s, incubated for 10 min at 4°C and centrifuged for 10 min at 14,000 rpm and 4 °C. The supernatants were removed and evaporated to dryness using speed vacuum concentrator (SpeedVac) and stored at −80°C until analysis. QCs were prepared by pooling all the tissue integrated in the study. Extraction was done using similar protocol use for the samples. Supernatant were aliquots in 34 tubes considering similar quantity. Then they were treated like samples.

Untargeted metabolomics approach applied in this study has been described in our previous paper ^57^. Metabolome profile of fat tissue was obtained from 264 samples including 32 QCs and 232 adipose tissue samples from mice v-AT and sc-AT. Data was acquired from both positive and negative polarities. Data acquisition in positive mode has been finalized in two days of continues run, while negative mode completed in three separated analytical runs. Raw data were transformed to mzXML format using MSConvert (Proteo Wizard 3.07155) and pre-processed for peak peaking, chromatogram alignment and isotope annotation using open access XCMS online (https://xcmsonline.scripps.edu). XCMS runs on UPLC-QExactive parameters by setting peak detection on Centwave. Overall, more than 20 thousands of M/Z values are sorted and aligned as the features of adipose tissue metabolome for the two acquisition polarities.

### Chemometrics and Pathway analysis

For analysis, raw data generated in targeted metabolomics were log2-transformed for each metabolite. Further normalization for across-batch signal drift was don using either QC-based model specifically *lowess*-model from open access web page (http://prime.psc.riken.jp/Metabolomics_Software/LOWESS-Normalization/) or non-QC based algorithms using “*dbnorm*” package. Notably, data for each mode of acquisition was treated separately for batch effect removal through either of QC-based or non QC-based model, and then merged for visual check and downstream differential analysis. A total of 239 different plasma metabolites detected in a human prospective cohort study study among which XANTHOSINE 5’-MONOPHOSPHATE was missing in QCs analyzed in positive modes.

Untargeted metabolomics data subjected to batch effect was also treated for normalization of across-batch signal drift using statistical methods implemented in the “*dbnorm*” package, which is also used for visualization of the outcome.

Metabolome signature in either of study, human study and animal experiment, was obtained by linear logistic regression model using lm (R ***Stats*** Package) and *limma* package (https://bioconductor.org), respectively. In the human cohort study, GFR is estimated by an equation developed by the Chronic Kidney Disease Epidemiology Collaboration (CKD-EPI) used as an outcome of renal impairment. CKD-EPI as estimator of kidney function calculated by gender and stratified by creatinine is assessed via following formula^58^ and recommended when eGFR values above 60 mL/min/1.73 m^2^.

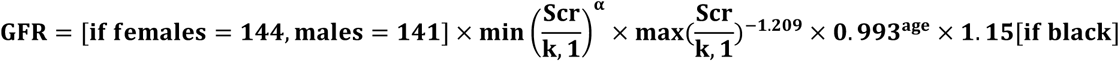

Where Scr is serum creatinine in mg/dl, k is factor (males=0.9, females=0.7) and α is-0.411 for males and,-0.329 for females, min indicates the minimum of Scr/κ or 1, and max indicates the maximum of Scr/κ or 1. Pearson’s r is calculated to rigorously estimate relationship between creatinine level measured in clinical routine and by MS. CIs at α < 0.05 (i.e. at 95% level) were produced upon transformation of pearson’s r to fisher’s z and calculated via following formula:

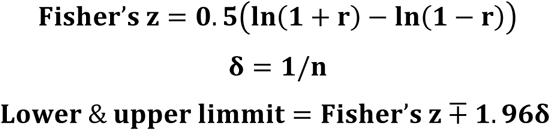

Subsequently, the association between CKD-EPI and plasma metabolome were investigated by multiple regression testing adjusted for age and sex.

In mouse model of obesity, we mainly focused on sc-AT metabolome signature driven by HFD after 8 weeks treatment, in comparison with their Ctrl counterpart. Subsequently, significant list was further searched and filtered against a Human Metabolome Database (HMDB; http://www.hmdb.ca) to keep only potential hits that were ultimately confirmed by MS2 spectra. Differential list in either of project, human prospective cohort study or animal experiment, was then subjected for over-representation analysis (ORA) using web interface of ConsensusPathDB (http://consensuspathdb.org) to pinpoint biochemical pathways that are dysregulated and may have a causative relationship to the phenotype. The list of identifiers was mapped to predefined KEGG pathways database enlisted by 4289 compound IDs. “p-value is calculated according to the hypergeometric test based on the number of physical entities present in both the predefined set and user-specified list of physical entities” (http://cpdb.molgen.mpg.de/). The selection criteria was that at least two metabolites representing the biological pathways displayed a significance level < 5% (q-value <0.05). Pathway visualization was done by BioCircos package (https://cran.r-project.org/web/packages/BioCircos).

## Acknowledgements

These research was supported by Swiss National Science Foundation [grant numbers: 33CM30-124087 and 310030_156771]

## References

1 Roerink, M. E., Bronkhorst, E. M. & van der Meer, J. W. Metabolome of chronic fatigue syndrome. Proc Natl Acad Sci U S A 114, E910, doi: 10.1073/pnas.1618447114 (2017).

2 Kurita, K. L., Glassey, E. & Linington, R. G. Integration of high-content screening and untargeted metabolomics for comprehensive functional annotation of natural product libraries. Proc Natl Acad Sci U S A 112, 11999–12004, doi: 10.1073/pnas.1507743112 (2015).

3 Davies, S. K. et al. Effect of sleep deprivation on the human metabolome. Proc Natl Acad Sci U S A 111, 10761–10766, doi: 10.1073/pnas.1402663111 (2014).

4 Newgard, C. B. Metabolomics and Metabolic Diseases: Where Do We Stand? Cell Metab 25, 43–56, doi: 10.1016/j.cmet.2016.09.018 (2017).

5 Sussulini, A. Erratum to: Chapters 1 and 11 of Metabolomics: From Fundamentals to Clinical Applications. Adv Exp Med Biol 965, E1–E2, doi: 10.1007/978-3-319-47656-8_14 (2017).

6 Dona, A. C., Coffey, S. & Figtree, G. Translational and emerging clinical applications of metabolomics in cardiovascular disease diagnosis and treatment. Eur J Prev Cardiol 23, 1578–1589, doi: 10.1177/2047487316645469 (2016).

7 Hocher, B. & Adamski, J. Metabolomics for clinical use and research in chronic kidney disease. Nat Rev Nephrol 13, 269–284, doi: 10.1038/nrneph.2017.30 (2017).

8 Long, J. Z. et al. Metabolomics annotates ABHD3 as a physiologic regulator of medium-chain phospholipids. Nature chemical biology 7, 763–765, doi: 10.1038/nchembio.659 (2011).

9 Subbaraj, A. K. et al. A large-scale metabolomics study to harness chemical diversity and explore biochemical mechanisms in ryegrass. Commun Biol 2, 87, doi: 10.1038/s42003-019-0289-6 (2019).

10 Tzoulaki, I., Ebbels, T. M., Valdes, A., Elliott, P. & Ioannidis, J. P. Design and analysis of metabolomics studies in epidemiologic research: a primer on -omic technologies. Am J Epidemiol 180, 129–139, doi: 10.1093/aje/kwu143 (2014).

11 Ala-Korpela, M. & Davey Smith, G. Metabolic profiling-multitude of technologies with great research potential, but (when) will translation emerge? Int J Epidemiol 45, 1311–1318, doi: 10.1093/ije/dyw305 (2016).

12 Fearnley, L. G. & Inouye, M. Metabolomics in epidemiology: from metabolite concentrations to integrative reaction networks. Int J Epidemiol 45, 1319–1328, doi: 10.1093/ije/dyw046 (2016).

13 Fuhrer, T. & Zamboni, N. High-throughput discovery metabolomics. Curr Opin Biotechnol 31, 73–78, doi: 10.1016/j.copbio.2014.08.006 (2015).

14 Nygaard, V., Rodland, E. A. & Hovig, E. Methods that remove batch effects while retaining group differences may lead to exaggerated confidence in downstream analyses. Biostatistics 17, 29–39, doi: 10.1093/biostatistics/kxv027 (2016).

15 Stein, C. K. et al. Removing batch effects from purified plasma cell gene expression microarrays with modified ComBat. BMC Bioinformatics 16, 63, doi: 10.1186/s12859-015-0478-3 (2015).

16 Reisetter, A. C. et al. Mixture model normalization for non-targeted gas chromatography/mass spectrometry metabolomics data. BMC Bioinformatics 18, 84, doi: 10.1186/s12859-017-1501-7 (2017).

17 Fernandez-Albert, F. et al. Intensity drift removal in LC/MS metabolomics by common variance compensation. Bioinformatics 30, 2899–2905, doi: 10.1093/bioinformatics/btu423 (2014).

18 Reese, S. E. et al. A new statistic for identifying batch effects in high-throughput genomic data that uses guided principal component analysis. Bioinformatics 29, 2877–2883, doi: 10.1093/bioinformatics/btt480 (2013).

19 Dunn, W. B. et al. Procedures for large-scale metabolic profiling of serum and plasma using gas chromatography and liquid chromatography coupled to mass spectrometry. Nature protocols 6, 1060–1083, doi: 10.1038/nprot.2011.335 (2011).

20 Llorach, R., Urpi-Sarda, M., Jauregui, O., Monagas, M. & Andres-Lacueva, C. An LC-MS-based metabolomics approach for exploring urinary metabolome modifications after cocoa consumption. Journal of proteome research 8, 5060–5068, doi: 10.1021/pr900470a (2009).

21 Luan, H., Ji, F., Chen, Y. & Cai, Z. statTarget: A streamlined tool for signal drift correction and interpretations of quantitative mass spectrometry-based omics data. Analytica chimica acta 1036, 66–72, doi: 10.1016/j.aca.2018.08.002 (2018).

22 Kirwan, J. A., Broadhurst, D. I., Davidson, R. L. & Viant, M. R. Characterising and correcting batch variation in an automated direct infusion mass spectrometry (DIMS) metabolomics workflow. Analytical and bioanalytical chemistry 405, 5147–5157, doi: 10.1007/s00216-013-6856-7 (2013).

23 Rusilowicz, M., Dickinson, M., Charlton, A., O’Keefe, S. & Wilson, J. A batch correction method for liquid chromatography-mass spectrometry data that does not depend on quality control samples. Metabolomics : Official journal of the Metabolomic Society 12, 56, doi: 10.1007/s11306-016-0972-2 (2016).

24 Zelena, E. et al. Development of a robust and repeatable UPLC-MS method for the long-term metabolomic study of human serum. Analytical chemistry 81, 1357–1364, doi: 10.1021/ac8019366 (2009).

25 Bijlsma, S. et al. Large-scale human metabolomics studies: a strategy for data (pre-) processing and validation. Analytical chemistry 78, 567–574, doi: 10.1021/ac051495j (2006).

26 Kamleh, M. A., Ebbels, T. M., Spagou, K., Masson, P. & Want, E. J. Optimizing the use of quality control samples for signal drift correction in large-scale urine metabolic profiling studies. Analytical chemistry 84, 2670–2677, doi: 10.1021/ac202733q (2012).

27 van der Kloet, F. M., Bobeldijk, I., Verheij, E. R. & Jellema, R. H. Analytical error reduction using single point calibration for accurate and precise metabolomic phenotyping. Journal of proteome research 8, 5132–5141, doi: 10.1021/pr900499r (2009).

28 Eilers, P. H. A perfect smoother. Analytical chemistry 75, 3631–3636 (2003).

29 Cleveland, W. S., Kleiner, B. & Warner, J. L. Robust statistical methods and photochemical air pollution data. J Air Pollut Control Assoc 26, 36–38 (1976).

30 Wang, S. Y., Kuo, C. H. & Tseng, Y. J. Batch Normalizer: a fast total abundance regression calibration method to simultaneously adjust batch and injection order effects in liquid chromatography/time-of-flight mass spectrometry-based metabolomics data and comparison with current calibration methods. Analytical chemistry 85, 1037–1046, doi: 10.1021/ac302877x (2013).

31 Wehrens, R. et al. Improved batch correction in untargeted MS-based metabolomics. Metabolomics : Official journal of the Metabolomic Society 12, 88, doi: 10.1007/s11306-016-1015-8 (2016).

32 Broadhurst, D. et al. Guidelines and considerations for the use of system suitability and quality control samples in mass spectrometry assays applied in untargeted clinical metabolomic studies. Metabolomics : Official journal of the Metabolomic Society 14, 72, doi: 10.1007/s11306-018-1367-3 (2018).

33 van Baar, B. L. et al. IS addition in bioanalysis of DBS: results from the EBF DBS-microsampling consortium. Bioanalysis 5, 2137–2145, doi: 10.4155/bio.13.172 (2013).

34 Stokvis, E., Rosing, H. & Beijnen, J. H. Stable isotopically labeled internal standards in quantitative bioanalysis using liquid chromatography/mass spectrometry: necessity or not? Rapid communications in mass spectrometry : RCM 19, 401–407, doi: 10.1002/rcm.1790 (2005).

35 Redestig, H. et al. Compensation for systematic cross-contribution improves normalization of mass spectrometry based metabolomics data. Analytical chemistry 81, 7974–7980, doi: 10.1021/ac901143w (2009).

36 Salerno, S., Jr. et al. RRmix: A method for simultaneous batch effect correction and analysis of metabolomics data in the absence of internal standards. PloS one 12, e0179530, doi: 10.1371/journal.pone.0179530 (2017).

37 Sysi-Aho, M., Katajamaa, M., Yetukuri, L. & Oresic, M. Normalization method for metabolomics data using optimal selection of multiple internal standards. BMC Bioinformatics 8, 93, doi: 10.1186/1471-2105-8-93 (2007).

38 Jacob, L., Gagnon-Bartsch, J. A. & Speed, T. P. Correcting gene expression data when neither the unwanted variation nor the factor of interest are observed. Biostatistics 17, 16–28, doi: 10.1093/biostatistics/kxv026 (2016).

39 De Livera, A. M. et al. Statistical methods for handling unwanted variation in metabolomics data. Analytical chemistry 87, 3606–3615, doi: 10.1021/ac502439y (2015).

40 Shen, X. T. et al. Normalization and integration of large-scale metabolomics data using support vector regression. Metabolomics : Official journal of the Metabolomic Society 12, doi: ARTN 89 10.1007/s11306-016-1026-5 (2016).

41 Ritchie, M. E. et al. limma powers differential expression analyses for RNA-sequencing and microarray studies. Nucleic acids research 43, e47, doi: 10.1093/nar/gkv007 (2015).

42 Chen, C. et al. Removing batch effects in analysis of expression microarray data: an evaluation of six batch adjustment methods. PloS one 6, e17238, doi: 10.1371/journal.pone.0017238 (2011).

43 Wen, B., Mei, Z., Zeng, C. & Liu, S. metaX: a flexible and comprehensive software for processing metabolomics data. BMC Bioinformatics 18, 183, doi: 10.1186/s12859-017-1579-y (2017).

44 Fernandez-Albert, F., Llorach, R., Andres-Lacueva, C. & Perera, A. An R package to analyse LC/MS metabolomic data: MAIT (Metabolite Automatic Identification Toolkit). Bioinformatics 30, 1937–1939, doi: 10.1093/bioinformatics/btu136 (2014).

45 Johnson, W. E., Li, C. & Rabinovic, A. Adjusting batch effects in microarray expression data using empirical Bayes methods. Biostatistics 8, 118–127, doi: 10.1093/biostatistics/kxj037 (2007).

46 Giordan, M. A Two-Stage Procedure for the Removal of Batch Effects in Microarray Studies. Stat Biosci 6, 73–84, doi: 10.1007/s12561-013-9081-1 (2014).

47 Kimura, T. et al. Identification of biomarkers for development of end-stage kidney disease in chronic kidney disease by metabolomic profiling. Sci Rep 6, 26138, doi: 10.1038/srep26138 (2016).

48 Pong, S. et al. 12-hour versus 24-hour creatinine clearance in critically ill pediatric patients. Pediatr Res 58, 83–88, doi: 10.1203/01.PDR.0000156225.93691.4F (2005).

49 Leoncini, G. et al. Creatinine clearance and signs of end-organ damage in primary hypertension. J Hum Hypertens 18, 511–516, doi: 10.1038/sj.jhh.1001689 (2004).

50 Chen, J. et al. Metabolomics Reveals Effect of Zishen Jiangtang Pill, a Chinese Herbal Product on High-Fat Diet-Induced Type 2 Diabetes Mellitus in Mice. Front Pharmacol 10, 256, doi: 10.3389/fphar.2019.00256 (2019).

51 Adam, O., Wolfram, G. & Zollner, N. Effect of alpha-linolenic acid in the human diet on linoleic acid metabolism and prostaglandin biosynthesis. Journal of lipid research 27, 421–426 (1986).

52 Kuchel, O. G. & Shigetomi, S. Dopaminergic abnormalities in hypertension associated with moderate renal insufficiency. Hypertension 23, I240–245, doi: 10.1161/01.hyp.23.1_suppl.i240 (1994).

53 Ponte, B. et al. Reference values and factors associated with renal resistive index in a family-based population study. Hypertension 63, 136–142, doi: 10.1161/HYPERTENSIONAHA.113.02321 (2014).

54 Guessous, I. et al. Associations of ambulatory blood pressure with urinary caffeine and caffeine metabolite excretions. Hypertension 65, 691–696, doi: 10.1161/HYPERTENSIONAHA.114.04512 (2015).

55 Ackermann, D. et al. CYP17A1 Enzyme Activity Is Linked to Ambulatory Blood Pressure in a Family-Based Population Study. Am J Hypertens 29, 484–493, doi: 10.1093/ajh/hpv138 (2016).

56 Caputo, T. et al.: Systemic approaches reveal anti-adipogenic signals at the onset of obesity– relatedinflammation in white adipose tissue. Cellular and Molecular Life Sciences (under revision)

57 Kowalczuk, L. et al. Proteome and Metabolome of Subretinal Fluid in Central Serous Chorioretinopathy and Rhegmatogenous Retinal Detachment: A Pilot Case Study. Transl Vis Sci Technol 7, 3, doi: 10.1167/tvst.7.1.3 (2018).

58 Burballa, C. et al. MDRD or CKD-EPI for glomerular filtration rate estimation in living kidney donors. Nefrologia 38, 207–212, doi: 10.1016/j.nefro.2017.02.007 (2018).

