## Supplementary Figures 1,2 for "Visualization and normalization of drift effect across batches in metabolome-wide association studies"

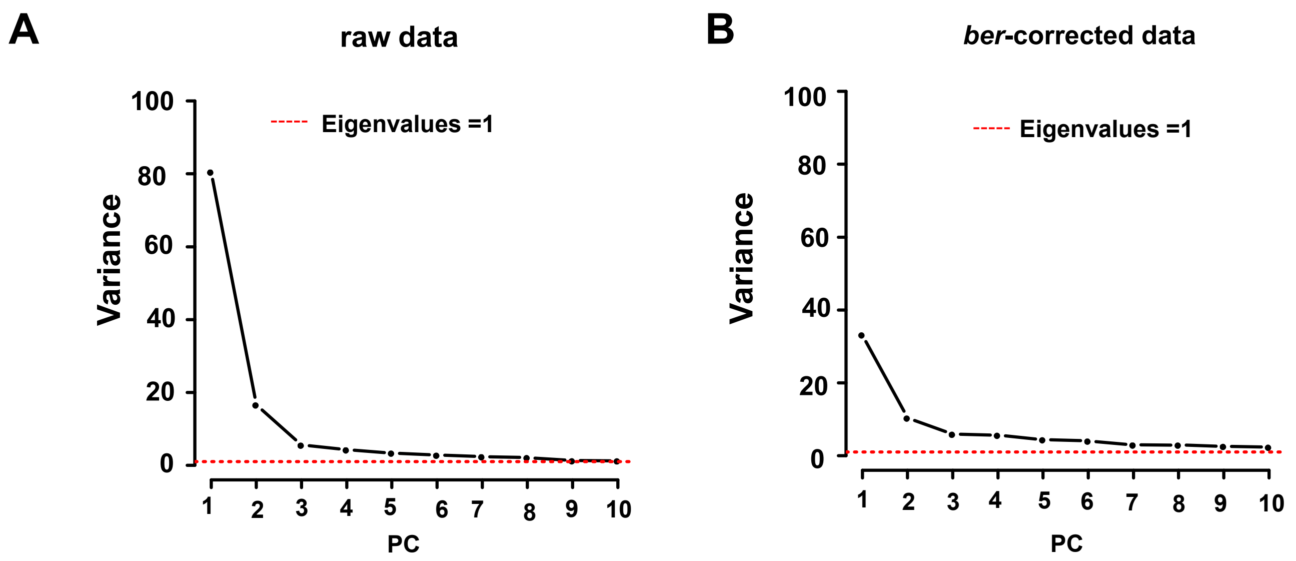


**Supplementary Figure 1.** Principal components analysis (PCA) of plasma metabolome in QC samples of the cohort study. Scree plots of 10 Principal Components (PCs) show that in raw data the three first principle components explains about 80 % of variation among the QC samples (**A**) while this number decreased to less than 40 % in the data adjusted for signal drift using *ber*-statistical model (**B**). Red dashed line colored shows the Kaiser criterion (eigenvalues higher than 1)

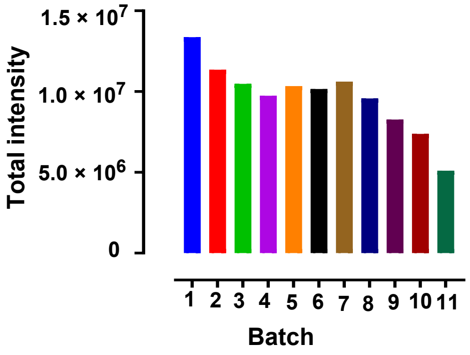


**Supplementary Figure 2.** Influence of batch order on total signal in the plasma metabolome in QC samples of the human cohort study. Total Ion Current (TIC) for each batch are reported for raw data showing over-time drift in signal intensity for identical QC samples.
