## Supplementary Figure 3 for "Visualization and normalization of drift effect across batches in metabolome-wide association studies"

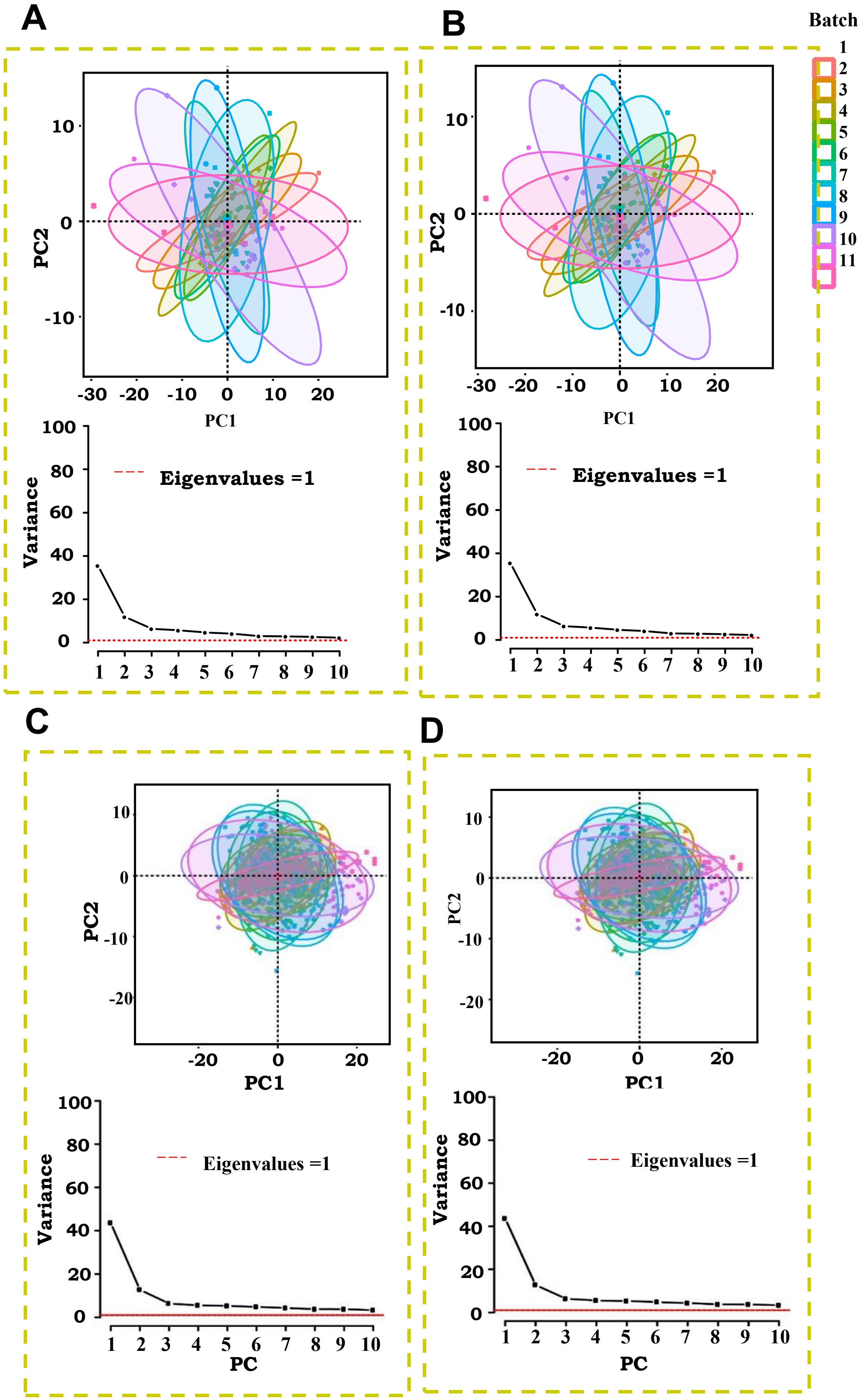


**Supplementary Figure 3.** Detection of batch effect and its correction in LC-MS targeted metabolomics analysis of the human cohort study. Principal Component Analysis (PCA) score plots show the absence of a strong source of variation associated to inter analytical runs in after correction of data for batch effect via parametric *ComBat* model in both QCs (**A**) and 1079 study samples (**C**) with less than 50% of variance explained by the first three PCs demonstrated by scree plot in the lower panels. Similar results were obtained for the accommodation of batch effect by applying *non-parametric ComBat* model in QCs (**B**) and 1079 study samples (**D**)
