## Supplementary Figure 4 for "Visualization and normalization of drift effect across batches in metabolome-wide association studies"

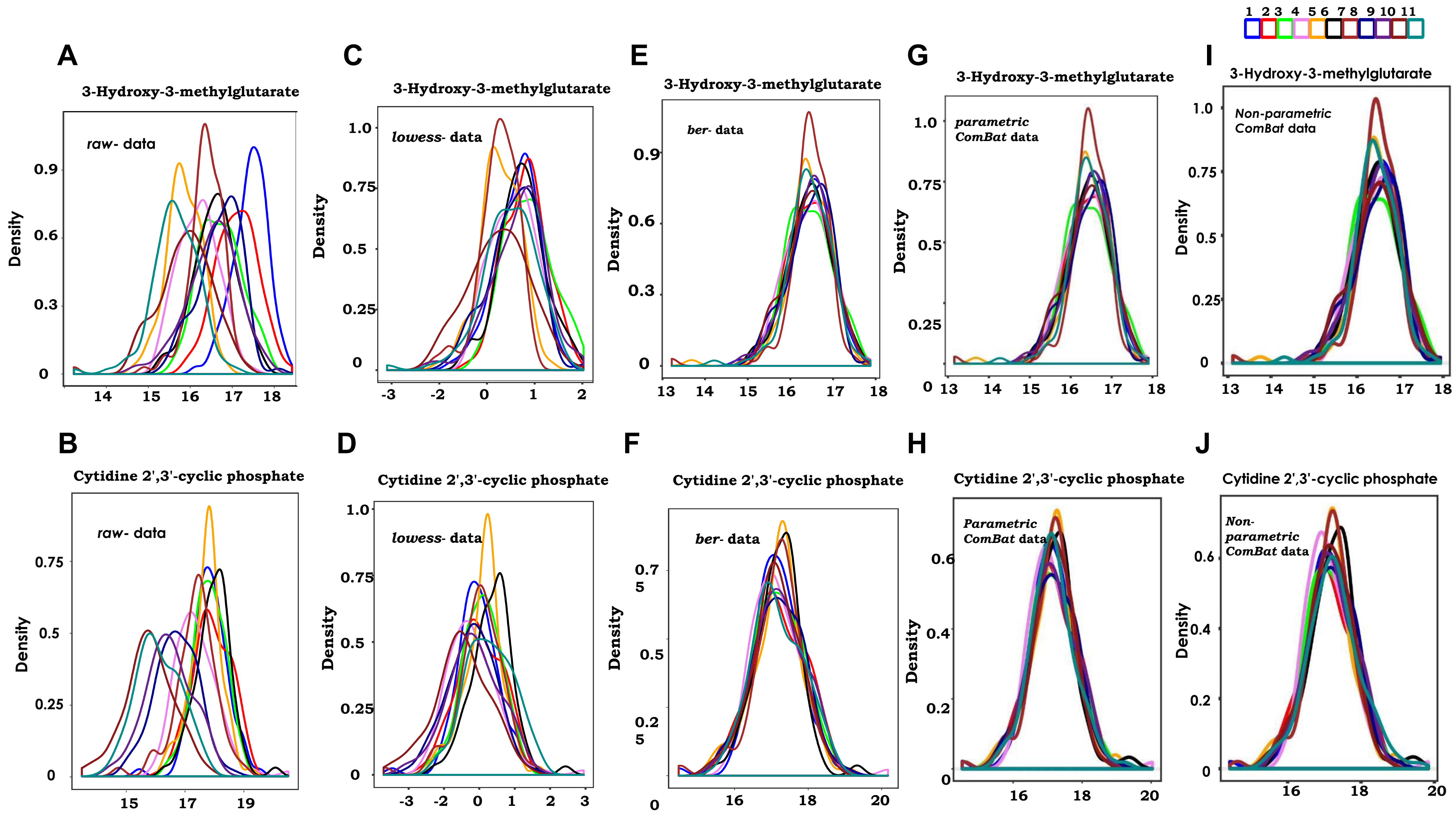


**Supplementary Figure 4.** Investigation of batch effect in LC-MS targeted metabolomics analysis of 1079 samples of a human study at the level of individual features. Probability density function (PDF) plots of 3-Hydroxy-3-methylglutarate and Cytidine 2',3'-cyclic phosphate across different experimental runs in the human cross-sectional study analyzed in 11 batches showed shift in the distribution of signals in *raw*- data (**A,B**). After data adjustment with either *lowess*- (**C,D**), *ber*- (**E,F**), parametric (**G,H**), or non parametric (**I,J**) *ComBat,* the average distribution of pdfs of these metabolites better overlap across batches. 3-Hydroxy-3-methylglutarate and Cytidine 2',3'-cyclic phosphate peaks are detected in positive and negative mode of acquisitions, respectively.
