## Supplementary Figure 5 for "Visualization and normalization of drift effect across batches in metabolome-wide association studies"

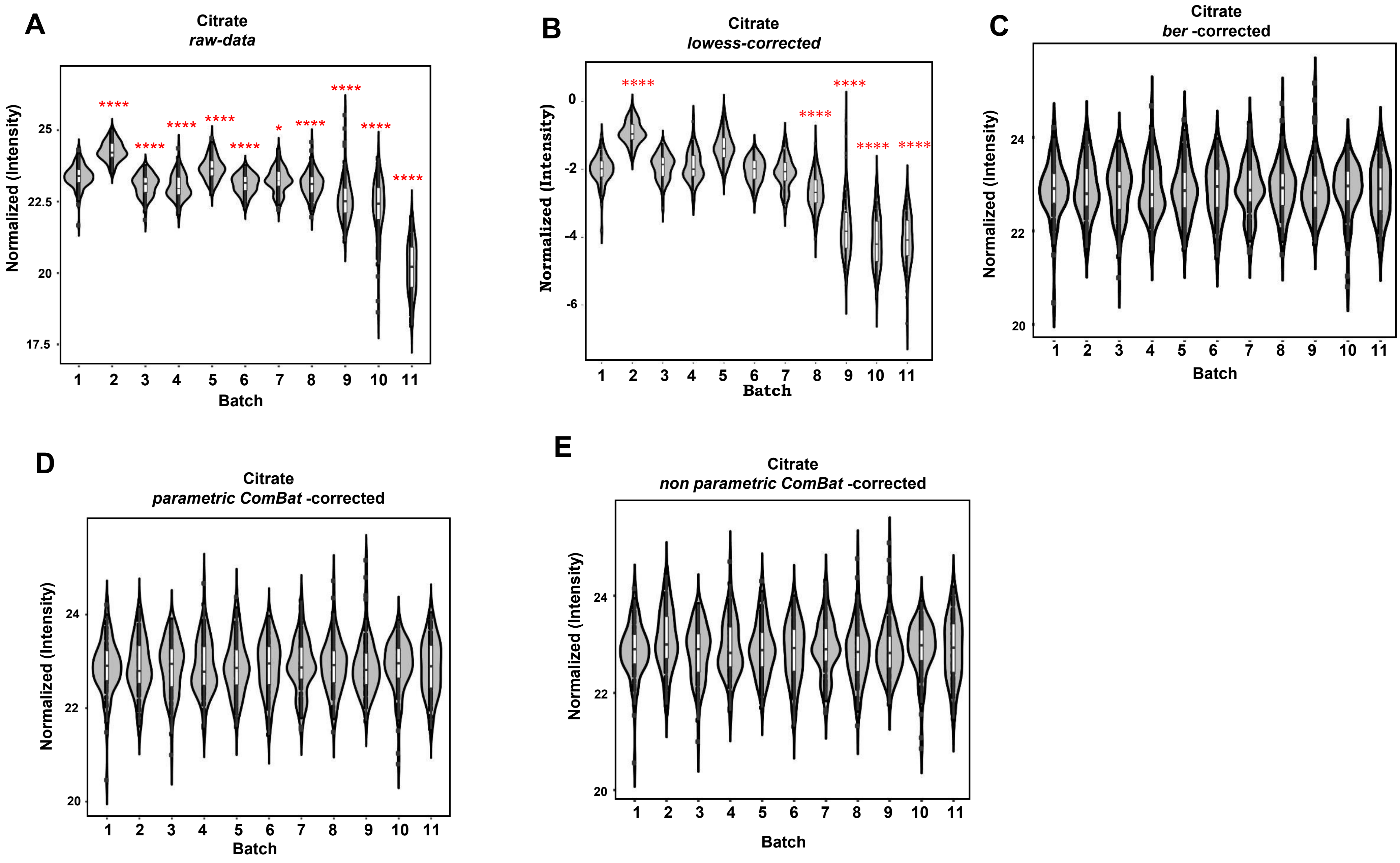


**Supplementary Figure 5.** Profile of citrate across sample sets in different dataset in Human cohort study. Violin plots of superposed scatter plots across batches for citrate showed the shift of average distribution across-batch in in raw data (A) which still presented in some batches in *lowess*-adjusted data (**B**). In contrast, data corrected with *ber* (**C**), parametric *ComBat* (**D**) and non parametric *ComBat* (**E**) are successfully corrected for the inter batches signal shift of citrate levels. Dots correspond to the level of citrate in each sample of the cross-sectional study and the average citrate level in each batch is define by boxplot. Two-tailed Student's *t*-test was used to calculate the significant changes of citrate mean level in different batches compared to the batch 1. *, **, *** and **** indicate p-value <0.05, <0.01, <0.001, <0.0001, respectively.
