## Supplementary Figures 6,7 for "Visualization and normalization of drift effect across batches in metabolome-wide association studies"

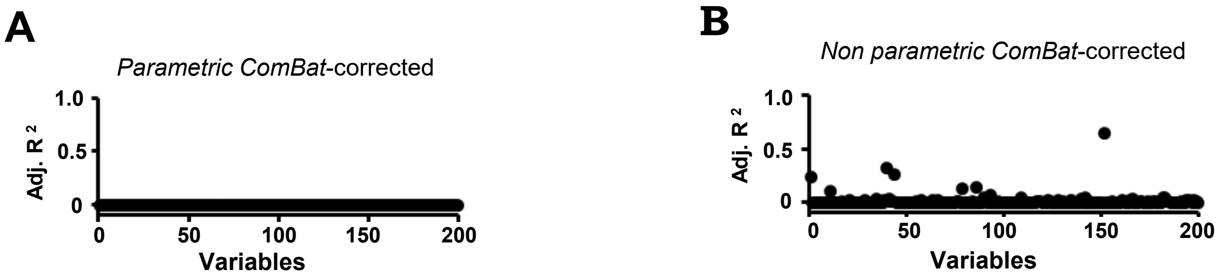


**Supplementary Figure 6.** Detection of batch effect and its correction in LC-MS targeted metabolomics analysis of 1079 samples of the cohort study . Adjusted Coefficient of Determination (Adjusted R-squared) estimated by regression model in data normalized with Parametric (**A**) and non parametric *ComBat* (**B**). A higher dependency of variability to the batch level is detected in non parametric compared to parametric *ComBat*-corrected data, as demonstrated by a variability closer to zero for all the detected features (variables) in panel A.

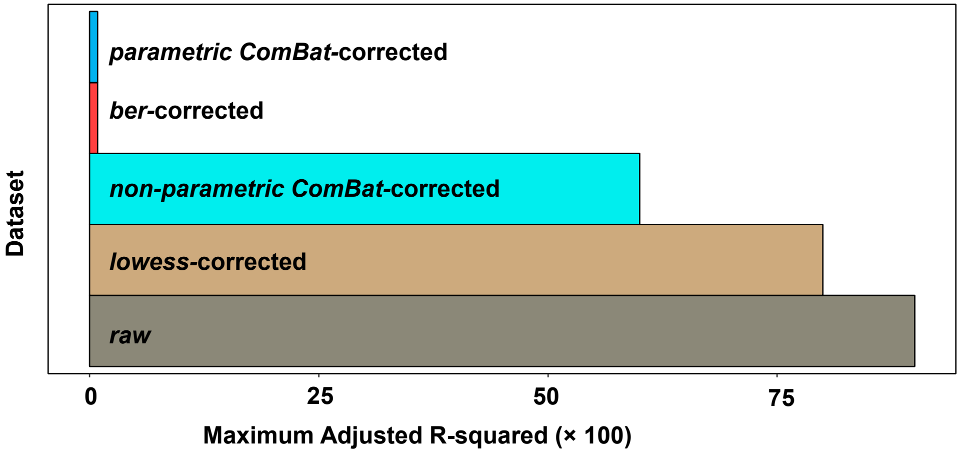


**Supplementary Figure 7.** Score performance calculation for normalization of batch effect drift. Adjusted Coefficient of Determination (Adjusted R-squared) is estimated by regression model. The maximum absolute Adjusted R-squared is calculated for each dataset and presented as bar plot. Colors represent datasets either non corrected, or corrected with different models. In the human cross-sectional study study, among the corrected dataset, lowess owns the highest maximum detected variability.
