## Supplementary Figure 8 for "Visualization and normalization of drift effect across batches in metabolome-wide association studies"

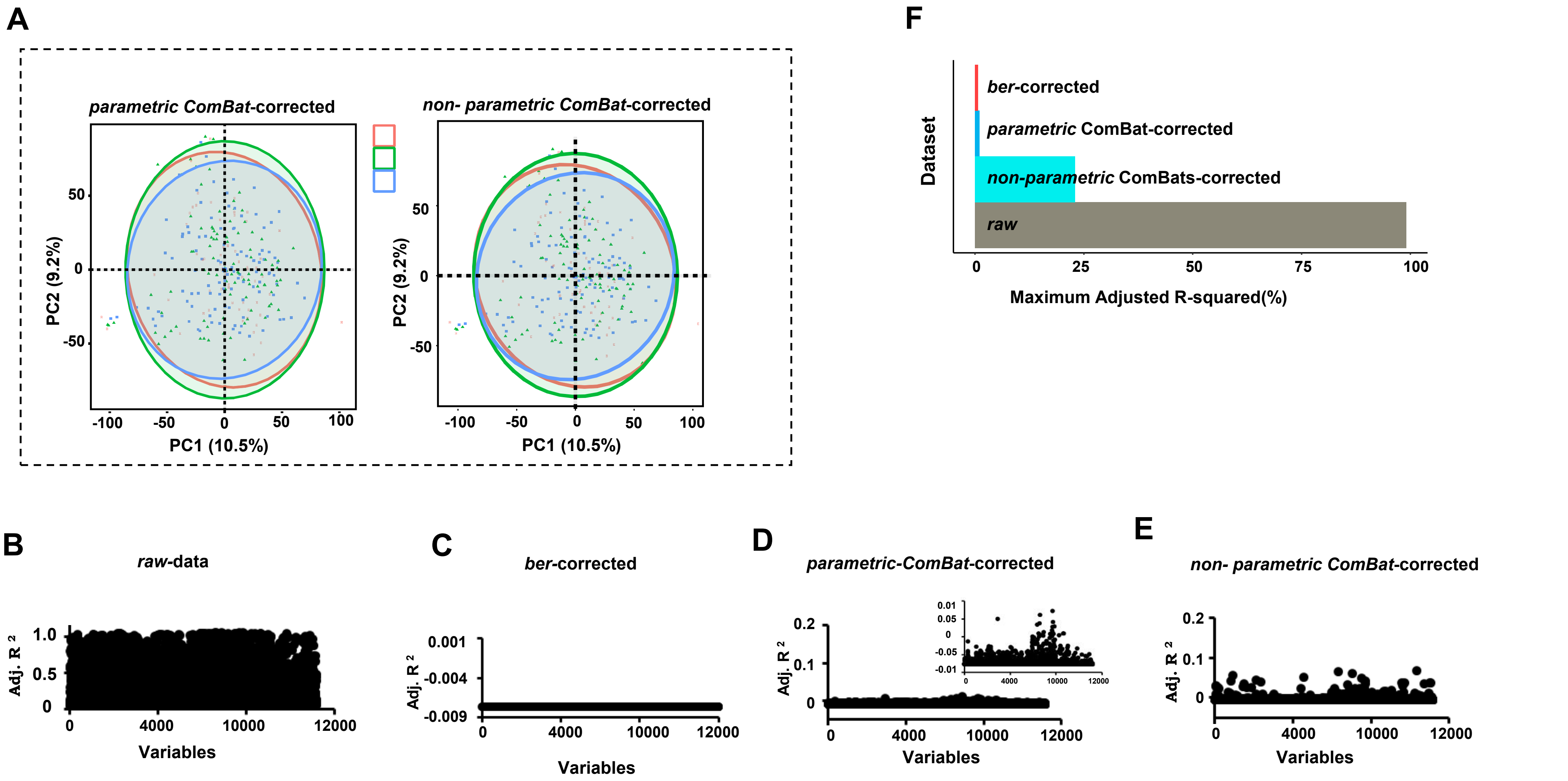


**Supplementary Figure 8.** Graphical check for the performance of statistical models in batch effect removal in LC-MS untargeted metabolomics analysis of 264 adipose tissue samples. **(A)** Principal Component Analysis (PCA) score plot of parametric and non parametric *ComBat*-corrected data do not show any separated clusters of samples along the experimental runs. Adjusted Coefficient of Determination (Adjusted R-squared), as estimated by regression model, shows that the variability dependent to the batch level in raw data **(B)** is about 100%, while a profound decrease to almost zero is detected in the *ber*-corrected dataset **(C)**. Reduced association between feature variability and batch level is also observed in parametric *ComBat*-corrected dataset, as visualized in both auto-scaled and adjusted scaled graph **(D)**, and in non parametric *ComBat*-corrected datasets **(E)**. The negative Adjusted R-squared values detected here are usually determinant of poor fitted model to assess variability. The score plot **(F)** summarizes the maximum dependency estimated in each dataset.
